# Integrated control of non-motor and motor efforts during decision between actions

**DOI:** 10.1101/2023.02.03.526983

**Authors:** Élise Leroy, Éric Koun, David Thura

## Abstract

Humans daily life is characterized by a succession of voluntary actions. Since energy resources are limited, the ability to invest the appropriate amount of effort for selecting and executing these actions is a hallmark of adapted behavior. Recent studies indicate that decisions and actions share important principles, including the exchange of temporal resources when the context requires it. In the present study, we test the hypothesis that the management of energy resources is shared between decision and action too. Healthy human subjects performed a perceptual decision task where they had to choose between two levels of effort to invest in making the decision, and report it with a reaching movement. Crucially, motor difficulty gradually increased from trial to trial depending on participants’ decision performance. Results indicate a relatively mild impact of the increasing motor difficulty on the choice of the non-motor (decision) effort to invest in each trial and on decision performance. By contrast, motor performance strongly decreased depending on both the motor and decisional difficulties. Together, the results support the hypothesis of an integrated management of energy resources between decision and action. They also suggest that in the context of the present task, the mutualized resources are primarily allocated to the decision-making process to the detriment of movements.

## INTRODUCTION

Human daily behavior is characterized by a succession of decisions ultimately expressed by movements. This requires the expenditure of energy resources, whose amount vary depending on the difficulty of the task and on the effort that one is willing to invest in carrying out this interactive behavior ^1^. The notion that effort is costly is supported by extensive experimental data. For example, activities requiring effort increase the response of the sympathetic nervous system, particularly in relation to blood pressure and pupil dilation, and induce the release of norepinephrine ^2^. As a result, individuals usually tend to avoid cognitive or motor effort when possible (but see ^3,4^). In other words, if a task offers the same amount of reward but imposes different levels of effort to obtain it, subjects typically choose the option associated with the minimum level of effort ^5–7^. Importantly, the willingness of individuals to exert effort during an activity decreases with the amount of effort already invested in this activity ^8^. This indicates that the energy resources necessary for the production of a costly behavior are limited, and that the choice of the level of effort to invest in the decision and in the action is crucial to guarantee an adapted and effective behavior.

Although decisions are always ultimately expressed via actions, cognitive and motor efforts are most often studied separately from each other. Recent behavioral studies, including ours, indicate however that decision and action are closely linked, sharing important principles and showing a high level of integration during goal-directed behavior ^9–19^. For instance, human subjects decide faster and with less precision in order to focus on their actions when the motor context in which a choice is made is demanding ^16^. Similarly, when the temporal cost of a movement is larger than usual, humans can shorten the duration of their decisions to limit the impact of these time-consuming movements ^18^. Conversely, if the sensory information guiding the choice is weak and the decision takes time, humans and monkeys shorten the duration of the movements expressing this choice ^12,17,20^. Individuals thus seem capable of sharing temporal resources, movement time for decision time, and vice versa, in order to determine a global behavior duration rather than optimizing the durations of decisions and actions separately. This mechanism is conducive to reward rate optimization ^21–23^.

The present study aims to test a complementary aspect of this hypothesis of an integrated control of decision and action. We propose that during decision between actions, the management of the effort-related energy resources is also integrated at the decision and action level in order to insure proficient behavioral performances. Such integrated control can take several forms, leading to different predictions. For instance, a simple yet intuitive possibility is that available energy resources are equitably allocated between decision and action depending on the respective effort context in which the behavior takes place. In such case, choosing to devote a large amount of effort on a decision will impact the performance of movements executed to express this choice and, conversely, if the effort required to perform an accurate movement is increased, the choice to engage in a difficult decision and the performance on that decision should decrease (figure 1). Alternatively, if decision and action effort-related energy resources are managed independently from each other, one should observe weak interactions between variations of decisional and motor difficulties and subjects’ decisional and motor performances (figure 1).

**Figure 1:**
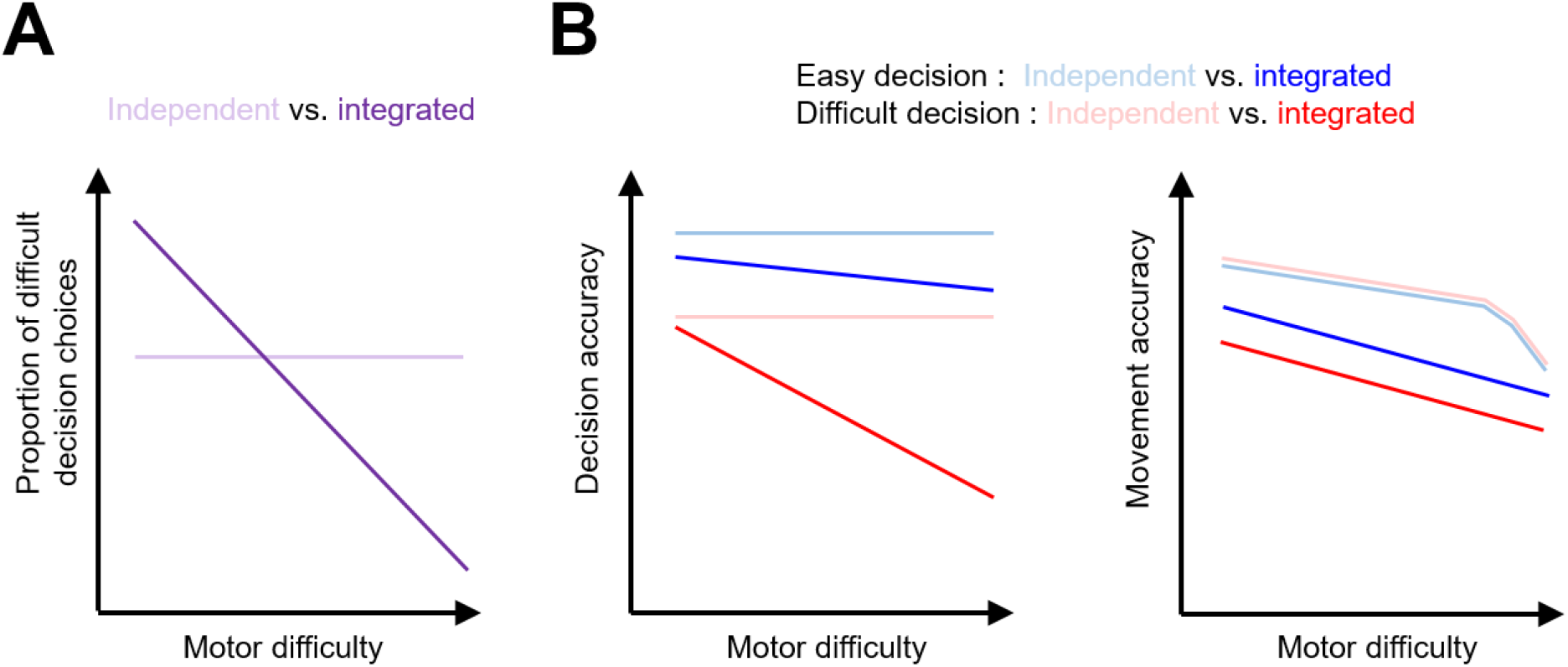
Predictions about the behavioral effects of an independent or integrated management of the decisional and motor effort-related energy resources. *A*. An independent management of resources predicts that the choice to engage in a difficult decision should not vary as a function of the effort required to perform an accurate movement. Alternatively, an integrated management of resources predicts that the choice to engage in a difficult decision will decrease if the effort required to perform an accurate movement increases. *B*. An independent management of resources predicts that decision performance should not vary depending on motor difficulty, regardless of the decision difficulty, easy (blue) or difficult (red). Similarly, motor performance should be only mildly impacted by an increased motor difficulty, because increasing resources can be allocated to the motor process when needed. In case of an integrated management of resources however, decisional performances should decrease if motor difficulty increases, especially for difficult decisions. Additionally, motor performance should be impacted by both motor and decisional difficulty.

## RESULTS

Thirty-two healthy human participants performed a new behavioral paradigm (figure 2) during a single experimental session. The goal of the subjects was to accumulate a total of 200 points to complete the session. To earn points, they had to choose at the beginning of each trial the amount of effort they wanted to invest in making a perceptual decision: either an effortful decision, potentially earning 5 points if correct, or an easy decision, earning only 1 point if correct. After making that choice, they had to make the corresponding perceptual decision and report it by executing an arm movement toward a visual target. Crucially and unknown to the subjects, the size of the movement targets was linearly and inversely indexed to the number of accumulated points during the session, progressively increasing the required motor control during the session. Importantly too, the points (5 or 1) that subjects chose to engage at the beginning of the trial were lost in case of a perceptual decision error, but not in case of an inaccurate movement, i.e. if they failed to reach the chosen target and stay in it within the required time windows. This task therefore allowed us to first observe the effect of the progressive increase of the motor accuracy requirement (or motor effort) on subjects’ choice of the non-motor effort to invest in a perceptual decision, and on their performance on that decisional process. Reciprocally, the task also allowed us to assess the effect of the perceptual decision difficulty on participants’ motor performance. Six additional participants performed the same procedure as the one described above except that the target size was smaller at the beginning of the session and did not evolve with the accumulation of points during the session. These subjects were tested to control that the reported effects were not due to fatigue or learning.

**Figure 2.**
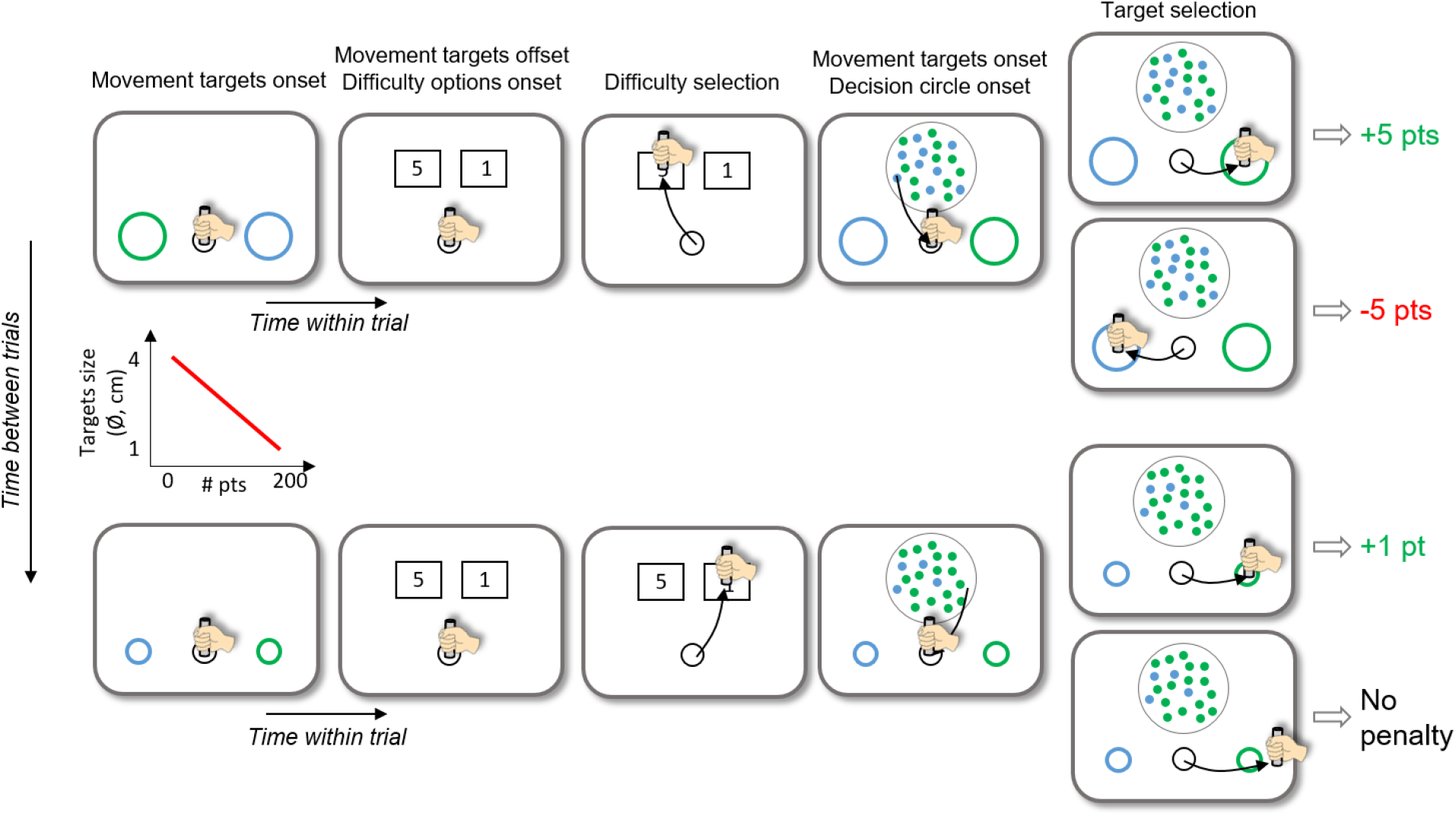
The top row illustrates the time course of a trial at the beginning of the session. Movement targets (a blue and a green circle) are first displayed to inform the subject about the accuracy requirement of the arm movement to execute later in the trial. The color of the targets at this stage is not informative of their color at the time of the perceptual decision. The diameter of the targets is 4cm during the first trial of the session. Difficulty options are then displayed. In this example the subject chooses “5”, which corresponds to a difficult (low coherence) perceptual decision to make. The decision circle containing 100 blue and green tokens, and the blue and green movement targets then appear. The dominant color among the tokens determines the correct target to select. The subject reports the decision by moving the handle in the target whose color corresponds to her/his choice. The subject earns the amount of points she/he chose (“5” in this example) if she/he accurately reaches to the correct target. She/he loses the points if she/he accurately reaches the target corresponding to the wrong decision. After the first trial, the size of the movement targets evolves from trial to trial, being linearly and inversely indexed to the number of points accumulated during the session. As a consequence, at the end of the session (bottom row), when the subject gets close to 200 points, the target size is small (diameter close to 1cm) and the required motor control is high. As illustrated in this example, an integrated control of resources between decision and action predicts that subjects would choose in this situation an easy decision (“1”) more frequently than at the beginning of the session, when the require motor control was low. If the subject fails to reach or stop in the chosen target (whether correct or not), points are not deducted.

### General observations

Among subjects who experienced the reduction of target size with the accumulation of points (n=32), the median proportion of high effort choice during a session was 50%, with a large variability between subjects (min: 0%; max: 100%; SD = 35%, figure 3A). An integrated control of resources between decision and action predicted that subjects would adjust their choices of the effort to invest in the perceptual decision through the session, choosing the most difficult perceptual decision more frequently at the beginning of the session than at the end (as depicted in figure 2). This is because the motor control requirement is the lowest at the beginning of the session (movement targets being large) and the amount of points to earn to complete the session is high. However, we observed that out of 32 participants, 9 almost did not vary their effort choices through the session (6/32 subjects chose the easy option in more than 95% of the trials, 3/32 chose that option in less than 5% of trials).

**Figure 3.**
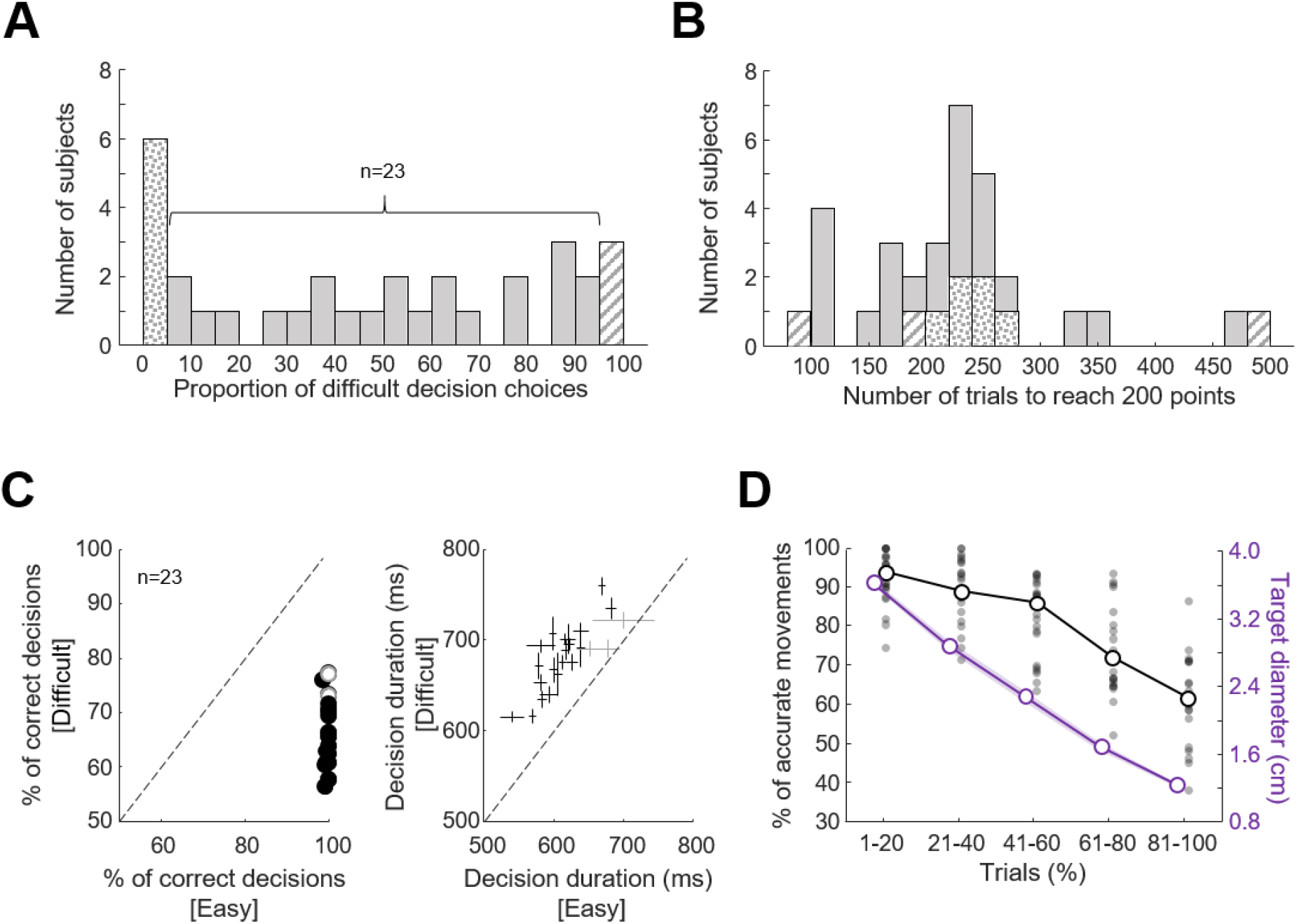
*A*. Distribution of the proportion of difficult perceptual decision choices (option “5”) among the 32 subjects who performed the main version of the task. The striped (dotted) bar highlights subjects who chose the difficult option more (less) than 95% of the trials. *B*. Distribution of the number of trials executed by the 32 subjects to earn 200 points and complete the session. Same convention as in A. *C*. Left panel: Comparison of subjects’ decision accuracy as a function of decision difficulty (Difficult: ordinate; Easy: abscissa). Circles illustrates individual subjects’ data. Black circles highlight subjects for which the difference between conditions is statistically significant (Chi-squared test, p<0.05). Right panel: Comparison of subjects’ decision duration as a function of decision difficulty (Difficult: ordinate; Easy: abscissa). Crosses illustrates individual subjects’ medians ± SD. Black crosses highlight subjects for which the difference between conditions is statistically significant (rank-sum test, p<0.05). *D*. Effect of the number of completed trials on subjects’ movement accuracy (black) and on target size (violet). Trials are sorted chronologically and a normalization is performed by grouping them in 5 quantiles. The open circles show median values for each quantile of trials across the population. The filled dots show individual subjects’ data for each quantile of trials.

The median number of trials to reach 200 points across the population was 227, with a large variability between subjects (min = 98; max = 481; SD = 88 trials, figure 3B). Subjects who did not adjust their effort choices during their session showed a particularly large variability in terms of the number of trials needed to complete the session (min = 98; max = 481; SD = 101 trials, figure 3B).

In the following analyses, we excluded the 9 subjects who systematically chose the same level of non-motor effort through their experimental session, as they were likely either insensitive (for the 3 subjects who chose the high effort option in more than 95% of the trials) or too sensitive (for the 6 subjects who chose the high effort option in less than 5% of the trials) to the decisional and/or motor difficulties manipulated in the experiment.

### Effect of decision difficulty on subjects’ decision behavior

We first verified that the two difficulty levels of the perceptual decision impacted the decision behavior of the 23 remaining subjects. To do so, we analyzed their decision duration and accuracy as a function of these two levels. As expected, we found that participants’ decision accuracy was usually lower when they made a difficult perceptual decision compared to when they had to make an easy one (medians: 65 versus 100%; Chi-square test for independence on the population: χ^2^ = 1124, p < 0.0001; Chi-square tests for independence on individual subjects, 21/23 with p < 0.05, figure 3C, left panel). Unsurprisingly too, subjects were overall slower to decide when faced with difficult perceptual decisions compared to when decisions were easy (medians: 662 versus 588ms, respectively; Wilcoxon rank-sum test on the population: Z=4.4, p < 0.0001; Wilcoxon rank-sum tests on individual subjects, 20/23 with p < 0.05, figure 3C, right panel). Given these results, we make the assumption in the following paragraphs that difficult decisions required the subjects to invest more non-motor effort compared to easy decisions.

### Effect of motor difficulty on subjects’ motor behavior

We then verified whether or not the motor accuracy requirement that increases with the number of accumulated points in this task impacted participants’ motor behavior. To do so, we analyzed their movement kinematics and accuracy as a function of the size of the targets. Because target size continuously varied from trial to trial, we normalized the number of trials performed by each subject by chronologically grouping them in 5 quantiles. As shown in figure 3D, the first 20% of trials were trials for which target size was the largest (because subjects’ scores were the lowest); Conversely, the last 20% of trials were the trials for which the target size was the smallest. As expected, the proportion of correct movements across the population significantly decreased depending on the number of trials performed during the session (Kruskal-Wallis test on the population, χ^2^ =67.1, p < 0.0001). There was also a trend for movement speed to decrease and duration to increase with the number of trials performed, but without reaching the level of significance (Kruskal-Wallis tests, χ^2^ = 5.6, p = 0.22; χ^2^ = 4.6, p = 0.33, respectively, supplementary figure 1, see also figure 5 for an analysis with trials grouped by decision difficulty). Given these results, we make the assumption in the following paragraphs that the smaller the target size, the more motor effort the subjects had to invest to execute accurate movements.

### Effect of increasing motor difficulty on decision behavior

Next, we investigated whether the increasing motor accuracy requirement (or motor effort) impacted the subjects’ willingness to invest effort in the perceptual decision-making. The prediction of an integrated control of decision and action-related energy resources was that with more motor effort, subjects would choose to make effortful perceptual decisions less frequently since they would have to devote more resources to face the more challenging actions (figure 2). However, we found at the group level that the proportion of difficult decision choices did not significantly vary depending on the level of motor difficulty (Kruskal-Wallis test, χ^2^ = 6.5, p = 0.16), despite the fact that a tendency for a decrease of that proportion with the increase of motor effort is visible (figure 4A). Indeed, at the individual level, we found that motor effort affected the proportion of difficulty choices in 15 out of 23 subjects (Chi-squared tests for independence, p < 0.05). Among them, the vast majority (12/15) overall decreased their proportion of high effort choices with the increase of motor effort. The duration of effort choices and the kinematics of movements directed to the effort options are shown for each effort option and against the session trials in supplementary figure 2.

**Figure 4.**
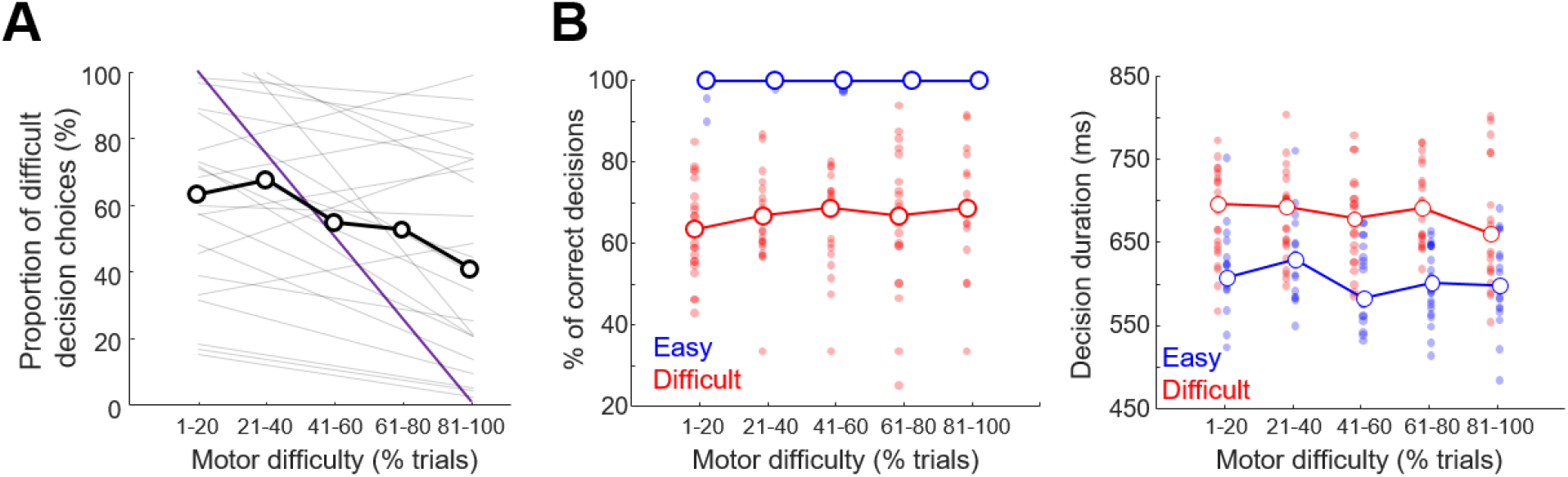
*A*. Proportion of difficult decision choices as a function of motor difficulty. As in figure 3D, trials are sorted chronologically and normalized by grouping them in 5 quantiles. Because target size strongly co-varies with the number of completed trials (figure 3D), trial number is a proxy of the motor accuracy requirement, and thus motor difficulty. Gray lines illustrate linear regressions through the data for each individual subject. The open dots show the median values for each trial quantile across the population. The violet line represents the hypothetical result of a perfectly shared management of resources between decisions and actions (figure 1): resources are initially only devoted to the decision part of the task because targets are big and movements easy; subjects thus only choose the difficult decision option; resources are linearly devoted to the movements as targets get smaller, and the proportion of difficult decision choices decreases; at the end of the session, resources are only devoted to movements because targets are small, subjects thus only choose the easy decision option to prioritize their invested efforts in executing accurate movements. *B*. Left panel: Proportion of correct perceptual decisions as a function of motor difficulty, with trials sorted as a function of decision difficulty (blue: easy; red: difficult). Right panel: Perceptual decision duration as a function of motor difficulty, with trials sorted as a function of decision difficulty (blue: easy; red: difficult). Same conventions as in figure 3D.

An integrated control of decision and action-related energy resources also predicted that with the increasing motor effort, the accuracy of the perceptual decisions would decrease and their duration would increase, especially for the most difficult ones. This is again because subjects would have to progressively devote more resources to face the increasingly challenging actions to execute to report these decisions, and consequently less resources would have been available to accurately make the perceptual decisions. Contrary to this prediction, we observed at the group level that difficult and easy decision performances were not affected by the increasing motor effort (Kruskal-Wallis test, χ^2^ = 6.27, p = 0.18; χ^2^ = 2.37, p = 0.67, respectively; figure 4B, left panel). Similarly, decision durations were not significantly impacted by the increasing motor difficulty through the session, regardless of the decision difficulty level (Kruskal-Wallis test, χ^2^ = 6, p = 0.2 for easy decisions; χ^2^ = 3.41, p = 0.49 for difficult decisions; figure 4B, right panel).

### Effect of the categorical decision difficulty on motor behavior

Finally, we analyzed the effect of the perceptual decision difficulty level on the way participants reported these decisions by reaching to the visual targets. To this aim, we analyzed the effect of motor difficulty on subjects’ movement accuracy, duration, speed and amplitude by grouping trials depending on the perceptual decision difficulty (figure 5). We observed more movement errors when participants’ reported a difficulty decision compared to when they expressed easy ones (ANCOVA, Difficulty: F=14.9, p = 0.0002). This effect did not depend on the size of the target, as no interaction between decision and action difficulties was observed (Difficulty x Trials: F=1.29, p=0.34). Interestingly, the effect is even more pronounced when only error decisions are included in the Difficult decision category (Difficulty x Trials: F=18.9, p<0.0001). We also observed a significant decrease of amplitude when movements followed difficult decisions compared to when they followed easy ones (Difficulty: F=12.2, p = 0.006), regardless of motor difficulty (Difficulty x Trials: F=0.23, p=0.63). Decision difficulty did not significantly impact movement speed (Difficulty: F=0.01, p = 0.95) nor duration (Difficulty: F=0.02, p = 0.89).

**Figure 5:**
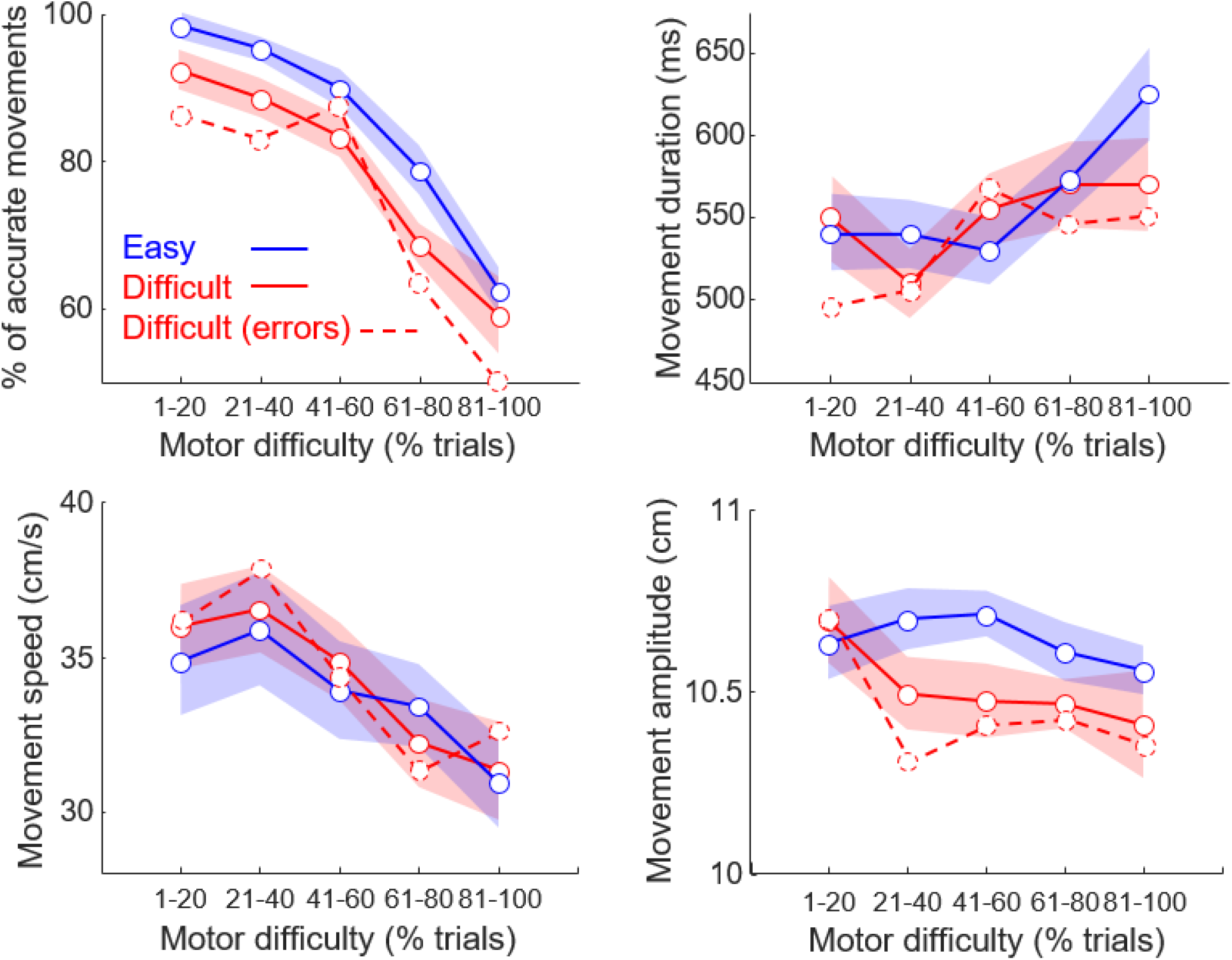
Effect of motor difficulty on subjects’ movement accuracy (top-left panel), duration (top-right), speed (bottom-left) and amplitude (bottom-right) with trials sorted according to the perceptual decision difficulty (red: difficult; blue: easy) and outcome (dotted red: difficult, wrong decisions). Same conventions as in Figure 4B, except that shaded areas illustrate the standard error around median values.

### Control subjects

To control that the effects reported above were not confounded by fatigue and/or learning, we describe in this last paragraph the behavior of 6 participants who performed the task in the exact same conditions as those described above, except that for them movement targets were smaller than those experienced by regular subjects at the beginning of the session and the size was kept constant during the session (i.e. it was not inversely and linearly related to the points accumulated during the session).

The median number of trials needed to reach 200 points across the six subjects was 194 ± 47, which is close to the median session duration experienced by subjects who performed the main experiment (227 ± 88 trials). The analysis of control subjects’ movement accuracy as a function of trials did not show any significant effect (Kruskal-Wallis test, χ^2^ = 2.46, p =0.65), suggesting that movement accuracy did not significantly suffer because of fatigue nor improved because of practice (supplementary figure 3A). Similarly, we found that the proportion of high effort choices did not significantly evolve as a function of session duration (Kruskal-Wallis test, χ^2^ = 2.13, p = 0.71, supplementary figure 3B). Interestingly, control subjects overall chose the high effort option less frequently at the beginning of their session compared to the 23 subjects who performed the main experiment. This makes sense in the light of an integrated management of resources between decision and action, as the size of the targets was smaller at session onset for the control subjects, facing them with more demanding motor control, probably discouraging them to choose the most effortful decision option.

Finally, we found that fatigue or learning did not impact the control subjects’ perceptual capacities, as their perceptual decision duration and accuracy did not significantly vary through the time course of the sessions (Kruskal-Wallis tests, χ^2^ = 4.6, p = 0.32; χ^2^ = 2.9, p = 0.57, supplementary figure 3C). Together, these analyzes on control subjects indicate that neither fatigue nor learning were the main factors explaining the results obtained on the 23 subjects who performed the main experiment.

## DISCUSSION

### Summary

In the present study, we asked healthy human subjects to choose the difficulty of perceptual decisions to make in individual trials, to make those decisions, and to report them with arm movements directed to visual targets in order to accumulate 200 points. Difficult decisions were worth 5 points, compared to only 1 point for easy ones. Crucially, the motor accuracy requirement increased with the accumulation of points. At the group level, we found that motor difficulty only mildly affected the proportion of difficult decisions chosen by participants, and had no significant impact on their decision duration and accuracy. By contrast, we found that motor difficulty strongly impacted movement accuracy, and that movement accuracy and amplitude were significantly reduced when a difficult decision was reported compared to when movements reported an easy one. Control analyses on additional subjects indicate a minor role of fatigue and/or learning in these effects.

The interaction between decisional and motor difficulties in the present work was designed to investigate the level of integration of the effort-related energy resource management between decision and action. According to the hypothesis of an integrated control of decision and action ^15,20,22^, resources are shared in a flexible and adapted way between these two processes, depending on the task demands. More specifically, an equitable distribution of resources predicts that increasing motor difficulty will force one to invest more effort in the motor process, leading to less frequent choices of the most difficult decision. It also predicts that performance while making these difficult decisions will decline with an increased motor effort. Alternatively, an independent management of the resources predicts that the proportion of difficult decision choices and decision performance will not vary depending on motor difficulty, and that movement accuracy and kinematics will not be influenced by decision difficulty (figure 1).

The present results do not fully support any of these two alternatives. Indeed, there is a trend for effort choices to be influenced by motor difficulty (figure 4A and supplemental figure 2B), but this influence is not as strong as expected if resources were equitably shared across decisions and actions. Moreover, the perceptual decision behavior appears very stable despite the increase of motor difficulty. If this argues at first sight for an essentially independent management of resources, the strong influence of decision difficulty on motor behavior (movement accuracy and amplitude) is not compatible with such independent management hypothesis.

One possible way to reconcile these results is to conceive that resources are shared between decision and action but not equitably, favoring in the present task the decision process over movements. In this view, subjects prioritized the allocation of their resources to the decision process, resulting in effort choices biased toward the difficult option despite the increase of motor difficulty, and, when a difficult decision was chosen, a maintenance of the decision accuracy figure 4A and 4B, left panel). Interestingly, this consistent accuracy is likely not the result of a simple speed-accuracy tradeoff that would have allowed subjects to compensate for less resources available for the decision by making longer decisions, resulting in constant accuracy^24^. Indeed, decision durations were overall stable within the time course of sessions too (figure 4B, right panel). A consequence of a prioritization of resources on the decision process is the “sacrifice” of the motor function. We indeed observed that movement accuracy was almost linearly reduced as a function of the increasing motor difficulty (figure 5, top-left panel). If resources were independently managed or equitably shared between decision and action depending on the task needs, we would have probably observed more stable movement performance through the sessions, at least until relatively late in these sessions. Finally, we observed that for a given target size, movements accuracy was lower and amplitude shorter when subjects reported difficult decisions compared to when they made easy ones, regardless of the size of the targets (figure 5, top-left and bottom right-panels). This suggests that the choice to allocate resources to make fast and accurate difficult decisions impaired participants’ ability to subsequently execute as accurate and ample movements as when they made easy decisions. Together, these results support the hypothesis of an integrated, but biased, management of the effort resources between decisions and actions, favoring in the present task decisions over actions.

The results discussed in the previous paragraphs indicate an important link between decision-making and motor control. Recent computational, behavioral, neurophysiological and clinical studies support this view, indicating that decision and action strongly influence each other^9,12,16–20,25–30^, operate according to the same ecologically-relevant principles^13,15,31^, share neural substrates^32–42^ and are often jointly altered in various neurological conditions^22,43^. For example, Thura and colleagues^12,17^ demonstrated in both monkeys and humans that when decision duration is long because of weak evidence, subjects shorten the duration of their movements to limit the loss of time on each trial and thus conserve their rate of reward at the session level. A similar interaction between decision and movement durations has been recently described in Parkinson’s patients^20^. Conversely, Reynaud and colleagues^16^ have shown that decisions are shortened and less likely to be correct when the motor context in which they are reported is demanding, requiring slow and accurate movements. The same authors then isolated the role of movement duration from effort in this effect, and showed that when the duration of the movement is lengthened, subjects shorten their decisions to limit the temporal devaluation of behavior^18^. Interestingly, the authors did not observe any consistent interaction between the decision and the action when the effort of the movement was manipulated. To explain this lack of effect, the authors proposed that unlike durations, effort-related energy costs were not as directly “exchangeable” between decisions and actions in the task they used. They also raised a key difference between effort and time, the fact that for a given behavioral success probability, effort is not necessarily always perceived as a cost (i.e. the effort “paradox”^3^) when time usually is^44–46^.

The present study validates both of these two explanations. Indeed, with a new behavioral task specifically designed to investigate the control of decision and motor-related energy resources, we observed that human subjects can exchange energy resources between decision and action depending on the task demands. This observation thus adds to our previous results showing that individuals are capable of sharing temporal resources in order to optimize their rate of succes^16–18^. The integrated control of effort-related resources described in the present report might be even more elaborated than a simple “dispatcher” of resources to each process considered in isolation. Indeed, this control seems to operate in a biased way, favoring in the present task decisions over actions. The most likely reason for such bias is that subjects strongly considered that decision outcomes had more task-goal implications than movement outcomes; a perceptual decision itself (i.e. regardless of the movement accuracy) allowing to earn or loose points whereas movement accuracy by itself was not rewarded nor penalized.

The present work also suggest that effort is probably not perceived as univocally penalizing across the population. Indeed, we often observed a large variability between the subjects, especially when we analyzed the choice of effort level to invest in the perceptual decision, both dependently and independently of the motor difficulty. As mentioned above, effort is generally felt to be aversive and difficult, which is why it tends to be avoided^5–7^. However, providing a lot of effort in a behavior can sometimes add value, and doing hard work can cause greater satisfaction than executing effortless tasks or even rest ^3,4^. Moreover, studies that investigated the impact of physical activity on cognitive abilities report that movements improve non-motor functions^47,48^. As a consequence, in some cases, or among some individuals, effort can be sought rather than avoided ^49^. This difference in value associated with effort may be one of the factors of variability we report between subjects.

A possible limitation of the study, related to the design of the task, concerns a possible learning-related familiarization with the decisional and motor difficulty experienced by the subjects through the time course of a session. However, several measures have been employed to limit this possibility (the training phase and the trial-to-trial variability of the decisional and motor difficulties) and the data obtained on 6 control subjects do not indicate a major impact of learning. The same is true for a potential role of motor and non-motor fatigue in this task. Data from control subjects do not indicate a decline in decisional and motor performance for a comparable length of experiment.

Another limitation of the study concerns the difficulty parameters of the decisions and actions which were the same for all subjects. As a consequence, difficulty levels and the resulting efforts were not necessarily perceived in the same way across the population. This probably explains part of the observed inter-subject variability, in particular the fact that 9 subjects did not change their proportion of difficult decision choices as a function of motor difficulty during their session. Another study using a staircase-type procedure to adapt the levels of difficulty to each subject could be more effective on this point.

Our results suggest a “sacrifice” of the motor system for the benefit of the cognitive system, possibly to prioritize the allocation of resources on the process allowing to earn or loose the points in the task. It would be interesting to assess whether or not the cognitive system can also sacrifice itself for the motor system. To this end, a complementary study in which difficulty parameters and task rules are switched between the decision and the action could be undertaken.

Finally, a distinction has been proposed between an account of effort based on computational or on metabolic costs^1^. Unlike physical effort, there does not appear to be a global metabolic cost for executing demanding non-motor tasks compared to automatic and effortless ones. In other words, the brain’s overall metabolic demands appear to change only mildly during engagement in non-motor behavior^1,50^. In the present task, motor difficulty was manipulated by means of varying the required level of movement accuracy, or movement control. This type of manipulation is probably different compared to a manipulation of load or resistance on the movements. It is thus possible that physical effort such as loaded or resistive movements induce more metabolic costs than motor control per se. By contrast, the cost of motor control is perhaps captured more accurately along the computational dimension, similar to that of perceptual decisions, which would have facilitated the integrated aspect of resource control between decisions and actions in our task. A very interesting question for future experiments is thus whether the present results are generalizable to other types of non-motor and motor efforts, tapping into different amounts of computational and metabolic costs.

## METHODS

### Participants

Thirty-height healthy human subjects (median age ± STD: 25 ± 4; 32 females; 35 right handed) participated in this study. All gave their consent before starting the experiment. The ethics committee of Inserm (IRB00003888, IORG0003254, FWA00005831) approved the protocol on June 7th 2022. Each participant was asked to perform one experimental session. They received a monetary compensation (10 euros per completed session) for participating in this study.

### Setup

The subjects sat in a comfortable armchair and made planar reaching movements using a handle held in their dominant hand. A digitizing tablet (GTCO CalComp) continuously recorded the handle horizontal and vertical positions (100 Hz with 0.013 cm accuracy). The behavioral task was implemented by means of LabVIEW 2018 (National Instruments, Austin, TX). Visual stimuli and handle position feedback (black cross) were projected by a DELL P2219H LCD monitor (60 Hz refresh rate) onto a half-silvered mirror suspended 26 cm above and parallel to the digitizer plane, creating the illusion that stimuli floated on the plane of the tablet.

### Behavioral task

Participants performed multiple trials of a multi-step decision-making task (figure 1). Each trial began with a small (Ø = 3cm) black circle (the starting circle) displayed at the bottom of the screen. To initiate a trial, the subject moved the handle in the starting circle and maintained the position for 300ms. Two colored circles (the movement targets: one blue, one green) were then displayed 180° apart of the starting circle for 200ms. The distance between the starting circle center and each movement target center was 10.9cm, with a trial-to-trial variability of 0.9cm. At this point subjects were informed about the accuracy requirement of their future movement (see how the size of the movement targets was determined below). The color of the targets at this stage is not informative of their color at the time of the perceptual decision.

Then the two movement targets disappeared and two rectangles appeared above the starting circle, separated from each other by 10 cm. In each rectangle a text informed the subject about the difficulty of the perceptual decision that she/he had to make in each trial: “1” for an easy decision, or “5” for a difficult decision. The subject had 1s to move the handle in the chosen rectangle and hold it for 500ms to validate this choice. She/he then returned to the starting circle and maintain the position for another 500ms to continue the trial.

Next, both rectangles disappeared and a large (Ø = 9cm) circle appeared on the screen (the decision circle). The decision circle was filled with 100 green and blue tokens, with different ratios between the two colors depending on the difficulty chosen at the beginning of the trial. “Difficult” decisions (“5”) were those in which the stimulus coherence (the ratio between the numbers of tokens of the two colors) was 53%, with a trial-to-trial variability of 2%; “Easy” decisions (“1”) were those in which the coherence was 75%, with a trial-to-trial variability of 2%. The subject task was to determine the dominant color in the decision circle, either blue or green. To express this perceptual decision, the participant moved the handle in the lateral target whose color corresponded to her/his choice and maintained this position for 500ms. The dominant color (blue or green) as well as the position of the green and blue movement targets relative to the starting circle were randomized from trial to trial. The maximum decision duration allowed (the time between the decision circle onset and movement onset) was 1s. The maximum movement duration allowed (the time between movement onset and offset) was 750ms.

At the end of the trial, a visual cue informed the subject about the outcome of the trial. The chosen target was surrounded by a green circle if she/he accurately reached the correct target, and by a red one if she/he accurately reached the wrong target. The subject earns the number of points corresponding to the chosen difficulty if the correct target was accurately reached. The goal of the subject was to earn a total of 200 points. In case of wrong decision (regardless of the accuracy of the movement), the number of points chosen at the beginning of the trial was subtracted. If the subject failed to reach or stop in the chosen target (inaccurate movement, whether it was the correct target or not), both movement targets turned orange and no points were deducted. To move on to the next trial, the subject moved the handle back in the starting circle and maintained the position for 500ms.

In the main experiment, performed by 32 out of 38 participants, the number of points accumulated by the subject determined the size of the movement targets. The diameter of these circles was set to 4 cm at the beginning of the session and it linearly decreased with the accumulation of points, reaching 1 cm at 200 points. As a consequence, the required motor control, and thus the motor difficulty, increased with the size reduction of the movement targets. We assumed that subjects increased their motor effort as movement targets get smaller with the number of trials performed and the number of points earned during the session. Six additional subjects performed the exact same task as the one described above except that the diameter of the movement targets was set to 2.5cm at the beginning of the session and was kept constant through the entire experiment. This control experiment was aimed to estimate effects that would not be a consequence of the increase of motor effort, such as fatigue or learning.

### Instructions provided to the subjects

To familiarize each participant with the task and with the manipulation of the lever on the tablet, a training phase was proposed prior to the experimental phase per se. During this training phase, subjects performed about 20 training trials where they could choose the difficulty of the decision to make (easy or difficult) and report these decisions by executing reaching movements to targets of 2.5 cm in diameter. The training phase was prolonged if subjects required so. During the experimentation phase, each subject was instructed to perform the task described above and they were informed that they needed to earn a total of 200 points to complete the session. Importantly, the 32 subjects who performed the main version of the task were not told about the decreasing size of the motor targets indexed to the accumulation of points. They were also not told about their number of points accumulated after each trial. We informed the subjects that there would be no scheduled breaks during the session, except in case of discomfort or real fatigue. No subject requested a break during their session.

### Data analysis and statistics

Data were collected by means of LabVIEW 2018 (National Instruments, Austin, TX), stored in a database (Microsoft SQL Server 2005, Redmond, WA), and analyzed off-line with custom-written MATLAB scripts (MathWorks, Natick, MA). Unless stated otherwise, data are reported as medians ± standard deviation.

Arm movement characteristics were assessed using the subjects’ movement kinematics. Horizontal and vertical arm position data (collected from the handle on the digitizing tablet) were first filtered using a tenth-degree polynomial filter and then differentiated to obtain a velocity profile. Onset and offset of movements were determined using a 3.75 cm/s velocity threshold. Peak velocity and amplitude was determined as the maximum value and the Euclidian distance between movement onset and offset, respectively.

An accurate movement is defined as a movement that reached a target (whether it is the correct target or not) and stayed in it for 500ms. In the main text of this report we only refer to movements executed to report the perceptual decisions. Kinematics of movements executed to select the difficulty of the decision at the beginning of the trial are illustrated in supplementary figure 2. Decision duration is defined as the time between the onset of the stimulus providing the visual evidence to the subject (the decision circle containing the 100 tokens) to the onset of the movement executed to report the decision. A decision is defined as correct if the correct target is chosen, regardless of the accuracy of the movement.

Chi-squared tests for independence were used to assess the effect of decision difficulty (easy or difficult) on individual subjects’ decision accuracy. Wilcoxon rank sum tests were used to assess the effect of decision difficulty on individual subjects’ decision duration. Chi-squared tests for independence were used to test the effect of motor difficulty, evaluated by chronologically grouping trials in 5 quantiles, on individual subjects’ movement accuracy and proportion of difficult choices. At the population level, Kruskal-Wallis tests were used to test the effect of motor difficulty on movement accuracy, decision accuracy, proportion of difficult choices, and on decision duration. Analyses of covariance (ANCOVAs) were used to assess the effect of decision difficulty, motor difficulty and their interaction on movement accuracy and kinematics (speed, duration, amplitude). The significance level of all statistical tests was set at 0.05, and highest levels of significance are reported when appropriate.

## Supporting information

https://www.dropbox.com/s/ffnjn84izswy5gr/LeroyKounThura_bioRxiv_SuppInfo_AsSub.pdf?dl=0

## Acknowledgements/Funding

This work is supported by a CNRS/Inserm ATIP/Avenir grant to David Thura. The authors thank Fadila Hadj-Bouziane and Clara Saleri Lunazzi for their valuable comments on the manuscript.

## AUTHORS’ CONTRIBUTION

ER, EK and DT designed the experiment

EK coded the task

ER collected the data

ER and DT conducted the analyses and prepared the figures

DT wrote the draft of the manuscript

ER, EK and DT revised the draft and approved the final version of the manuscript

## OPEN PRACTICES STATEMENT

This work’s data and codes are freely available upon request.

