## Supplementary material for "Integrated control of non-motor and motor efforts during decision between actions": https://www.dropbox.com/s/ffnjn84izswy5gr/LeroyKounThura_bioRxiv_SuppInfo_AsSub.pdf?dl=0

Lyon Neuroscience Research Center – ImpAct team
Inserm U1028 – CNRS UMR5292 – Lyon 1 University
16 avenue du Doyen Jean Lépine – 69676 Bron – France

*Corresponding author:*

David Thura

### Supplemental information

#### Supplementary figure 1

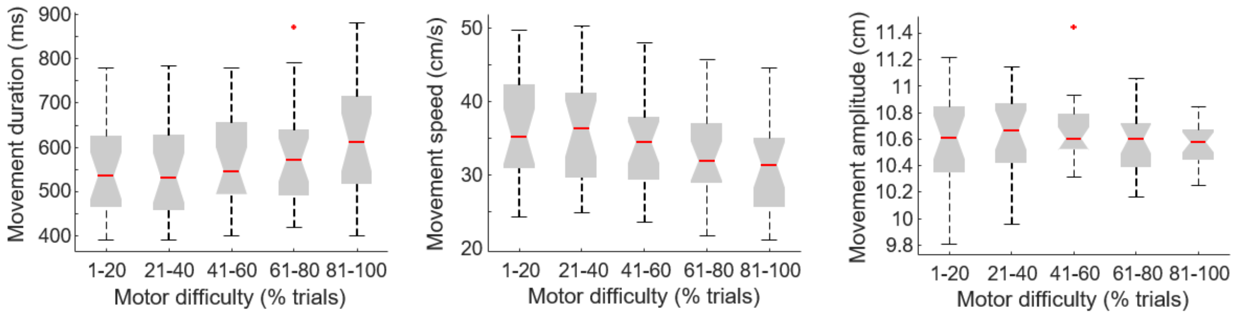


Effect of motor difficulty on subjects' movement duration (left), speed (center) and amplitude (right). Trials are sorted chronologically and a normalization is performed by grouping them in 5 quantiles. Because target size strongly co-varies with the number of completed trials (figure 2D in the main text), trial number is a proxy of the motor accuracy requirement, and thus of the motor difficulty. The box plots visually present the summary statistics (minimum, $1^{st}$ quartile, median, $3^{rd}$ quartile, maximum values as well as outliers (red crosses)) for each quantile of trials.

#### Supplementary figure 2

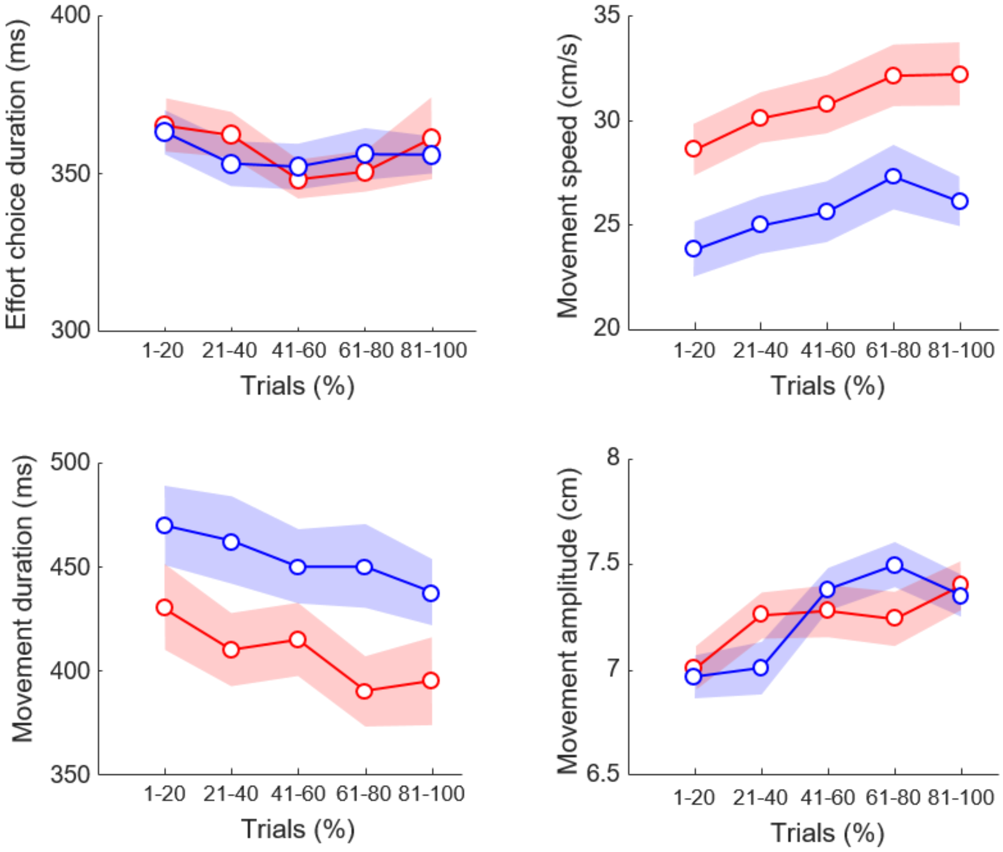


Top-left panel: Effort choice duration as a function of trials completed in the session, with trials sorted in each quantile as a function of the perceptual decision difficulty (blue: easy; red: difficult). Same conventions as in figure 5 in the main text. Bottom-Left: Duration of movements executed to select the level of difficulty of the perceptual decision. Movements were overall shorter when subjects selected the difficult option (red) compared to when they selected the easy option (blue); ANCOVA, Difficulty: F=10.3; p = 0.002. Top-right: Speed of movements executed to select the level of difficulty of the perceptual decision. Movements were overall faster when subjects selected the difficult option (red) compared to when they selected the easy option (blue); ANCOVA, Difficulty: F=11.1; p = 0.001. Bottom-right: Amplitude of movements executed to select the level of difficulty of the perceptual decision.

#### Supplementary figure 3

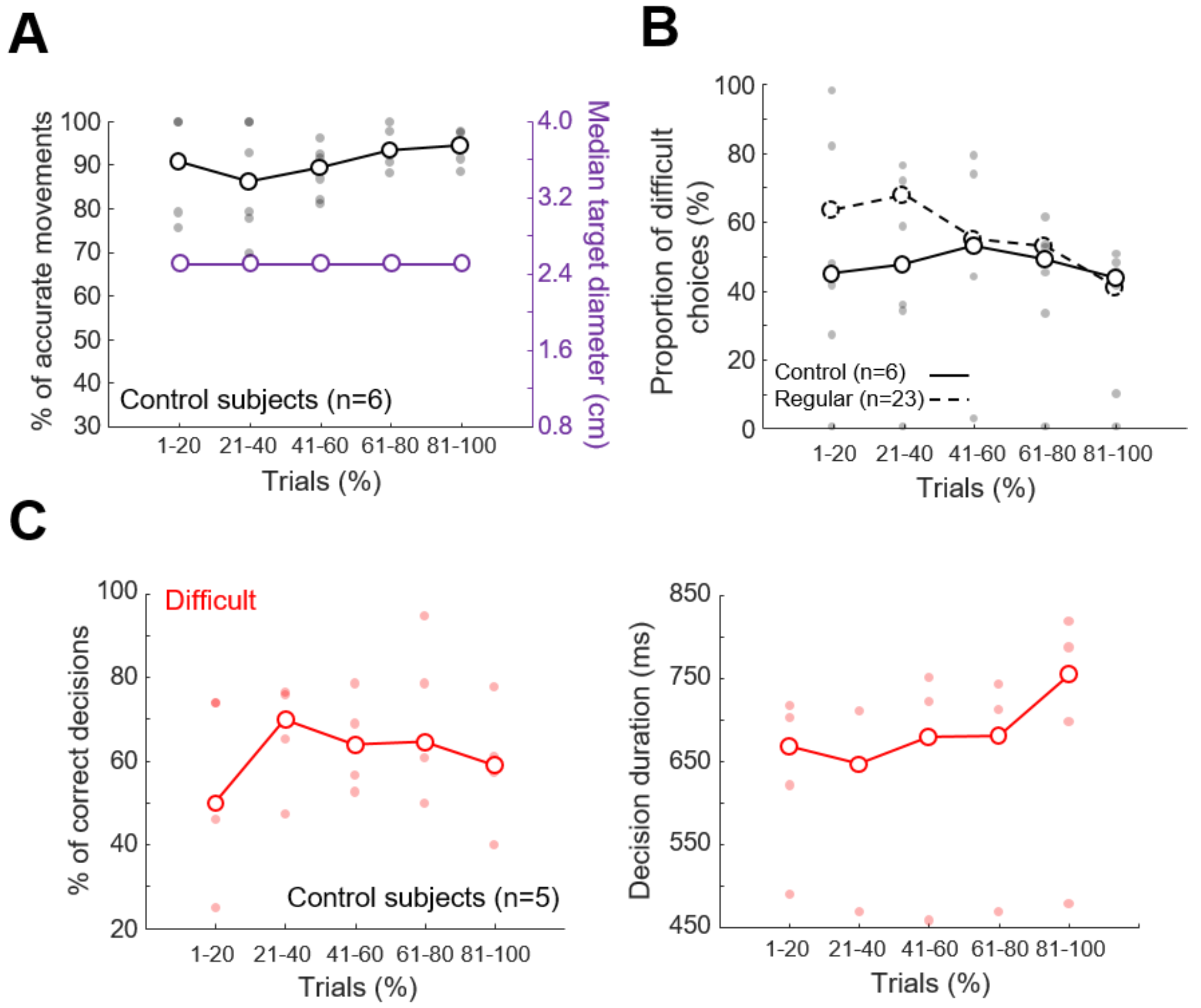


*A.* Effect of motor difficulty on control subjects' (n=6) proportion of accurate movements (black) and on target diameter (violet). Same conventions as in figure 3D in the main text. *B.* Effect of motor difficulty on control (n=6, solid black line) and non-control (n=23, dashed black line) subjects' proportion of difficult decision choices. The open circles show the median values for each trial quantile across each population of subjects. The filled black dots show individual control subjects' data for each quantile of trials. *C.* Left panel: Proportion of control subjects correct perceptual decisions as a function of motor difficulty, with trials restricted to those in which a difficult decision was chosen. Right panel: Same as left for perceptual decision duration. Same conventions as in B.
